# Hippocampal ERK2 dimerization is critical for memory reconsolidation and synaptic plasticity

**DOI:** 10.1101/2024.09.06.611755

**Authors:** Santiago Ojea Ramos, Candela Medina, María del Carmen Krawczyk, Julieta Millan, María Florencia Acutain, Arturo Gabriel Romano, María Verónica Baez, Francisco José Urbano, Mariano Martín Boccia, Mariana Feld

## Abstract

Extensive research has focused on extracellular-signal regulated kinase (ERK) 1/2 phosphorylation across various memory and plasticity models. However, the precise mechanisms linking ERK activity to memory stabilization and restabilization are still poorly understood, and the role of ERK1/2 dimerization remains unexplored. ERK dimerization is essential for the binding and activation of cytoplasmic targets, many of which are involved in memory and plasticity. In this study, we investigated the role of ERK2 dimerization in long-term memory and synaptic plasticity. We found that reactivation of a weak inhibitory avoidance (wIA) memory led to a significant reduction in hippocampal ERK2 dimerization. Furthermore, intrahippocampal infusion of DEL-22379 (DEL), an ERK dimerization inhibitor, following memory reactivation had a bidirectional effect: it blocked the reconsolidation of a strong inhibitory avoidance (sIA) memory but enhanced the reconsolidation of a wIA memory. Moreover, DEL administration blocked hippocampal ERK2 dimerization *in vivo* and impaired high-frequency stimulation-induced long-term potentiation (LTP) in hippocampal slices. These findings demonstrate that ERK2 dimerization occurs in the intact mouse nervous system and plays a pivotal role in plasticity and memory. While further research is needed, this study highlights the relevance of ERK dimerization in these processes.

## INTRODUCTION

The molecular mechanisms underlying learning and memory have been extensively studied using diverse experimental paradigms and techniques. Long-term plasticity is widely considered the cellular correlate of memory formation (Bliss and Collingridge, 1993; Bliss and Lømo, 1973). In line with this proposal, specific signaling pathways, such as extracellular signal-regulated kinase 1/2 (ERK1/2) have been implicated in both plasticity and memory processes (Atkins et al., 1998; Blum et al., 1999; English and Sweatt, 1997; Martin et al., 1997; Schafe et al., 2000; Walz et al., 1999). Pharmacological inhibition of ERK1/2 phosphorylation impairs both memory formation and neuronal plasticity (reviewed in Ojea Ramos et al., 2022).

ERK1/2 belongs to the mitogen-activated protein kinase (MAPK) family, a large group of Ser/Thr protein kinases originally associated with development, mitogenesis, and stress response (Lavoie et al., 2020). While ERK phosphorylation was initially thought to drive dimerization and subsequent nuclear translocation (Khokhlatchev et al., 1998), subsequent findings revealed that dimerization primarily mediates cytosolic function, enabling the activation of cytoplasmic targets via scaffolding proteins (Casar et al., 2008). Furthermore, the multiplicity of ERK1/2 cytosolic targets (Ahn, 2009; Earnest et al., 1996; Gong and Tang, 2006; Kelleher et al., 2004; Kneussel and Wagner, 2013; Schrader et al., 2006) supports the idea that extra-nuclear localization of the kinase is relevant for a wide variety of processes. DEL-22379 (DEL), a novel ERK dimerization inhibitor, effectively blocks hyperactive Ras-ERK1/2 signaling in tumor cells, impairing proliferation and transformation while circumventing resistance mechanisms previously described for phosphorylation inhibitors (Herrero et al., 2015).

In neuronal systems, ERK phosphorylation has been shown to regulate diverse processes, including long-term potentiation (Albert-Gascó et al., 2020), translation within specific subcellular compartments at active synapses (Pirbhoy et al., 2017), dendritic spine density in key brain areas involved in cognitive processes (Gao et al., 2011) and memory paradigms (Lavoie et al., 2020) and neurodegenerative pathologies (Albert-Gascó et al., 2020).

Stability of consolidated memories can be challenged by reactivation, opening a new period of sensitivity named “reconsolidation” and triggering molecular mechanisms involved in re-stabilization (Dudai and Eisenberg, 2004; Nader et al., 2000; Sara, 2000). Previous work from our group demonstrated that cytosolic ERK phosphorylation kinetics following inhibitory avoidance memory reactivation in mice regulates the reconsolidation, strength and persistence of the reactivated memory (Krawczyk et al., 2019, 2016, 2015), underscoring the importance of compartment-specific ERK activation in memory processes.

However, the role of ERK1/2 dimerization in synaptic plasticity and memory remains unknown. We hypothesized that ERK dimerization is a regulatory milestone for long-term memory (LTM) reconsolidation. Here we show for the first time that ERK2 dimerization can be pharmacologically inhibited in mice hippocampi without affecting its phosphorylation. Its inhibition blocked CA1 hippocampal LTP and affected memory reconsolidation of the inhibitory avoidance (IA) task, depending on the training strength. These findings shed light on a previously unrecognized role of ERK dimerization *in vivo*, emphasizing its importance in synaptic plasticity and memory.

## MATERIALS AND METHODS

### Animals

Different cohorts of CF-1 male mice aged either 5 weeks (weight range: 21 - 31 g, housed in groups of 3-5 individuals in 18 x 29 x 12 cm amber polysulfone cages at the Facultad de Ciencias Exactas y Naturales’ central animal facility) or 8-10 weeks old (weight range: 25 - 35 g, housed in groups of 18 - 22 individuals in stainless-steel 50 x 30 x 15 cm cages kept at the Laboratorio de Neurofarmacología de Procesos de Memoria, Facultad de Farmacia y Bioquímica) were used either for electrophysiological experiments or behavioral testing and biochemical analyses, respectively. Mice had *ad libitum* access to dry food and tap water and were maintained under a 12 h light/dark cycle (lights on at 6:00 AM) at 21 – 23 °C. Experiments were conducted during the light cycle. All procedures related to care and treatments of mice were approved by the local institutional animal care and use committees (CICUAL - CUDAP # 0044975-2016, Facultad de Farmacia y Bioquímica; CICUAL # 154b, Facultad de Ciencias Exactas y Naturales, Universidad de Buenos Aires, Argentina), and in compliance with the standard European Union and United States National Institutes of Health ethical guidelines for Care and Use of Laboratory Animals. All efforts were made to minimize any possible distress or suffering experienced by mice, as well as to reduce the number of animals required to generate reliable scientific data.

### Drugs

DEL-22379 (DEL), an ERK dimerization inhibitor (Herrero et al., 2015), was purchased from MedChemExpress (Cat. No.: HY-18932) and dissolved in dimethyl sulfoxide (DMSO) as vehicle (VEH). For behavioral experiments and hippocampal ERK2 dimerization assays, 0.5 μl of drug or VEH solution were injected per hemi-hippocampus (DEL final concentration: 400 nM). Epidermal growth-factor (EGF) was purchased from Thermo Fisher (Cat. No.: PHG0314) and diluted in sterile phosphate buffer saline (PBS) to a final concentration of 100 μM and 0.5 μL were injected per hemi-hippocampus.

### Hippocampal slices and extracellular field potential recordings

Electrophysiological recordings in slices were performed as described elsewhere (Rivero-Echeto et al., 2021). Briefly, mice were deeply anesthetized (ketamine 160 mg/kg, xylazine 8mg/kg) and decapitated. Coronal brain slices (300-400 μm thick) were obtained with a vibratome (PELCO, EasiSlicer, Ted Pella Inc., CA, USA), in chilled low-sodium/high-sucrose solution (250 mM sucrose, 2.5 mM KCl, 3 mM MgSO_4_, 0.1 mM CaCl_2_, 1.25 mM NaH_2_PO_4_, 0.4 mM ascorbic acid, 3 mM myo-inositol, 2 mM pyruvic acid, 25 mM D-glucose, and 25 mM NaHCO_3_), and kept at 35 °C for 30 min. Recovery chamber contained low Ca^2+^ / high Mg^2+^ normal artificial cerebrospinal fluid (ACSF; 125 mM NaCl, 2.5 mM KCl, 3 mM MgSO_4_, 0.1 mM CaCl_2_, 1.25 mM NaH_2_PO_4_, 0.4 mM ascorbic acid, 3 mM myo-inositol, 2 mM pyruvic acid, 25 mM d-glucose, and 25 mM NaHCO_3_ and aerated with 95% O_2_ / 5 % CO_2_, pH 7.4). Slices were perfused continuously with ACSF including bicuculline (20 μM; a GABA-A antagonist).

Borosilicate glass (2–3 MΩ) recording electrodes filled with ACSF + bicuculline solution were placed in the *stratum radiatum* area of the CA1 region of the hippocampus and field excitatory postsynaptic potentials (fEPSP) were recorded as measure of local excitatory synaptic transmission in the CA1 region. The stimulating electrode was placed at the Schaffer Collateral fibers and fEPSPs were evoked by test stimuli (0.033 Hz, 0.05-0.2 ms duration) every 30 sec. Long term potentiation (LTP) was induced by applying high frequency stimulation (HFS; 5 trains, 100 Hz, 10 pulses) after a baseline period of 10 min and recorded for 30 min. The magnitude of LTP was calculated as a percentage of the fEPSP slope value after HFS compared to baseline fEPSP mean slopes. Changes in LTP were compared in hippocampal slices incubated with either VEH or 1,33 μM DEL for 30 min. Signals were recorded using a MultiClamp 700 amplifier commanded by pCLAMP 10.0 software (Molecular Devices, CA, USA). Data was filtered at 5 kHz, digitized, and stored for off-line analysis.

### Intra-hippocampal injections

Hippocampal injections were performed under isoflurane anesthesia (induction: 3.5 %, maintenance: 1 %) using a stereotaxic apparatus (Stoelting) at the indicated time points at the following coordinates to target the dorsal hippocampus: AP: −1.90 mm from bregma, DV: −2.2 mm from the skull surface and +/-1.2 mm left and right from the midsagittal suture (Paxinos and Franklin, 2019) to bilaterally infuse the drugs. Injections were driven by an automatic pump (MD-1020, Bioanalytical Systems, Inc.) at a rate of 0.52 μL/min through a 30-gauge stainless steel needle attached to a 5 μl Hamilton syringe with PE-10 tubing. The drug infusion lasted 60 sec and the needle was left in place for another 30 sec to avoid reflux. Administered volume was 0.5 μl (either 400 nM DEL or VEH solution) per hemi-hippocampus. The incision was sutured and thoroughly disinfected. After the last testing session, animals were euthanized by cervical dislocation and brains were dissected, fixed by immersion in 4 % PFA overnight, and sectioned to verify the correct injection site placement. In all cases administration placement was confirmed in the target area (Suppl. Fig. 1).

### Inhibitory Avoidance task

Inhibitory avoidance behavior was assessed using a one-trial step-through protocol, which takes advantage of mice’s preference for dark and confined spaces, and has been previously described (Blake et al., 2008; Boccia et al., 2004). Two different shock intensities were used: a 0.3 mA - 3 sec shock for weak inhibitory avoidance induction (wIA), and a 0.4 mA - 3 sec shock for strong inhibitory avoidance (sIA) memory formation (Blake et al., 2014). Mice underwent the first retention test (RE) 48 h after training (TR), followed by a second retention test (TS) 24 h after RE. No shock was delivered during either test session. Previous work showed that RE also induces memory reactivation, and a non-reactivated (NR) group (animals that remained in the home cage) was used as a control (Boccia et al., 2007, 2010). Both retention tests were ended either when the mouse stepped into the dark compartment or failed to enter within 300 sec after the beginning of the test. After each retention test, mice were returned to their home cages. Step through latencies were manually registered using a manual chronometer.

### Sample preparation and ERK analysis

#### Neuronal cultures

Primary neuronal cultures were obtained and maintained as described before (Acutain et al., 2021; Kaech and Banker, 2006).

For the induction of ERK activation by EGF, 1 μl of the stock solution was diluted in 1 mL of culture medium and incubated in the culture for 5 min. Chemical LTP (cLTP) induction was achieved by a glycine stimulation based on a previously described (Swanger et al., 2013). After either glycine stimulation or control treatment for 3 min, neuronal cultures were cooled and homogenized by scraping cells in ice-cold lysis buffer (20 mM HEPES, pH 7.5, 10 mM ethylene glycol tetra-acetic acid, 40 mM glycerol 2-phosphate, 1 % (w/v) IGEPAL, 2.5 mM MgCl_2_, 2 mM sodium orthovanadate, 1 mM dithiothreitol, 0.5 mM PMSF, 50 mM Sodium Fluoride, 1 μg/ml Pepstatin A, 10 μg/mL Leupeptin, 10 μg/mL Aprotinin) as described previously (Pinto and Crespo, 2010). Samples were centrifuged at 13000 x g for 10 min at 4 °C, and the supernatants were collected and conserved at −20 °C.

### Hippocampal ERK activation and tissue protein extraction

For the study of hippocampal ERK activation by EGF, two consecutive injections were performed in each hemi-hippocampus: first either 0.5 μl of VEH (PBS) or 100 μM EGF followed by 0.5 μl of either VEH or 400 nM DEL. At the specified time points, mice were quickly euthanized by cervical dislocation, decapitated and the hippocampi were dissected. Each hemi-hippocampi was then homogenized using a Dounce tight homogenizer and 125 μL of ice-cold lysis buffer (20 mM HEPES, pH 7.5, 10 mM ethylene glycol tetraacetic acid (EGTA), 40 mM glycerol 2-phosphate, 1 % (w/v) IGEPAL, 2.5 mM MgCl_2_, 2 mM orthovanadate, 1 mM dithiothreitol, 0.5 mM PMSF, 50 mM Sodium Fluoride, 1 μg/ml Pepstatin A, 10 μg/mL Leupeptin, 10 μg/mL Aprotinin) by performing 8 strokes with a tight glass pestle. Samples were centrifuged at 13000 x g for 10 min at 4 °C, and the supernatants were collected and conserved at −20 °C.

### Native gel electrophoresis

Native polyacrylamide gel electrophoresis (PAGE) for ERK dimers detection was carried out as described previously (Pinto and Crespo, 2010). Protein levels in lysates were quantified by BCA Assay (Bio-Rad, USA) and 60 μg of protein were loaded into an 8 % bis-acrylamide gel and ran for 150 min at 70 V. Then samples were transferred to a nitrocellulose membrane (General Electric, USA) overnight at 125 mA and 4 °C.

### SDS-PAGE

ERK phosphorylation levels were assessed by means of denaturing gel electrophoresis as described before (Krawczyk et al., 2015). Briefly, 20 μg of protein were loaded in a 12.5 % bis-acrylamide gel and run for 90 min at 100 V. The samples were then transferred to a PVDF membrane (Thermo Scientific, USA) overnight at 300 mA and 4 °C and stored in TBS until probed with antibodies.

### Antibodies used and detection

Initially, membranes were blocked in 4% non-fat dry milk in TBS - 0.05 % Tween-20. A full list of antibodies used, and dilutions can be found in Table 1. Primary antibodies were incubated overnight at 4 °C and secondary antibodies were incubated for 1 h in TTBS at room temperature. ERK dimers were detected on nitrocellulose membranes by luminol chemiluminescence kit (Bio Rad ECL, USA) using X-Ray film (Agfa ORTHO CP-GU, cat. PTNAGCPGUM2, Voxel, Argentina). SDS-PAGE blots were developed using the same chemiluminescence kit in a GE Amersham Imager.

**Table 1.**
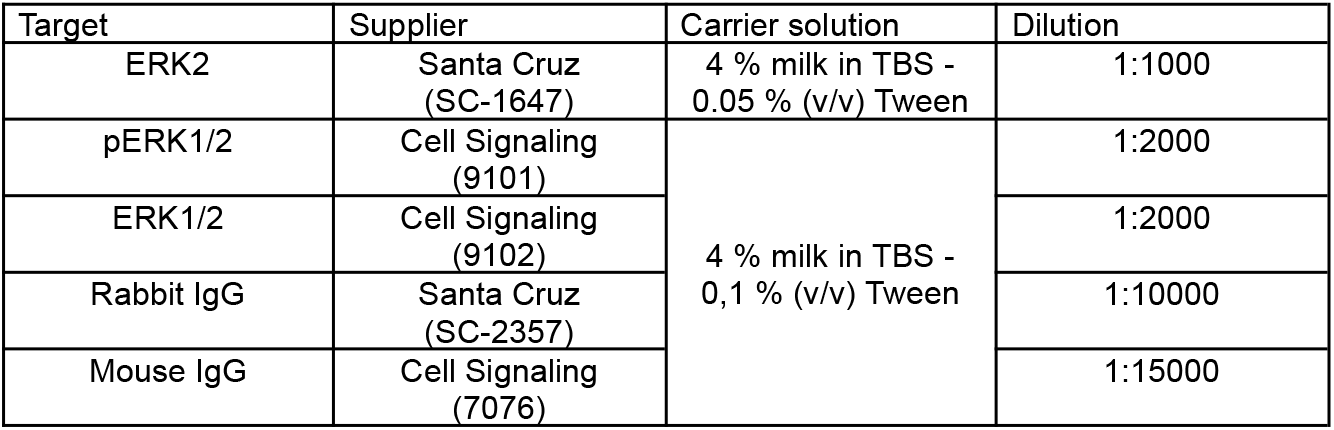

| Target | Supplier | Carrier solution | Dilution |
| --- | --- | --- | --- |
| ERK2 | Santa Cruz (SC-1647) | 4 % milk in TBS - 0.05 % (v/v) Tween | 1:1000 |
| pERK1/2 | Cell Signaling (9101) | 4 % milk in TBS - 0,1 % (v/v) Tween | 1:2000 |
| ERK1/2 | Cell Signaling (9102) |  | 1:2000 |
| Rabbit IgG | Santa Cruz (SC-2357) |  | 1:10000 |
| Mouse IgG | Cell Signaling (7076) |  | 1:15000 |

### Data analysis

Data analyses were conducted using R (version 4.2.2,83; R core team, 2021). Generalized linear models (GLMs) were employed using the glmmTMB package (version 1.1.6; Brooks et al., 2017). For behavioral data (latency to step-through), drug (VEH or DEL) and condition (RE or NR) were used as fixed factors. The model was fitted with a Gamma distribution and log link function. For western blot data, the model was fitted using a Gaussian distribution and identity link function. Repeated measures ANOVA with Time and Treatment as fixed factors was used to analyze the LTP experiment. The goodness-of-fit for each model was evaluated using the DHARMa package (version 0.4.6; Hartig, 2024). Visual inspection of residual plots did not reveal any concerning deviations from normality or homoscedasticity. ANOVA of the models is reported and, when applicable, post-hoc comparisons were performed using the emmeans package (version 1.8.5; Lenth, 2017). All reported p-values were adjusted for multiple comparisons with the Šidak correction method. Two-tailed Pearson coefficients were calculated to determine the correlation between phosphorylation and dimerization at each time point.

## RESULTS

### ERK dimerization in rodent nervous system and plasticity

The latency to enter the dark compartment in the inhibitory avoidance (IA) task partially depends on the intensity of the aversive stimulus delivered during training (Krawczyk et al., 2019). Previous work from our group showed that retrieval of a consolidated IA memory in CF-1 mice modulated hippocampal cytosolic phosphorylated ERK2 (pERK2) levels 45 min later, depending on the strength of the training stimulus. Weak IA (wIA) memory reactivation significantly increased pERK2, while strong IA (sIA) memory reactivation led to a significant decrease. Furthermore, intrahippocampal administration of the MEK1/2 inhibitor PD098059, 45 min after wIA retrieval strengthened memory at 24 h (Krawczyk et al., 2015).

Given the importance of cytosolic ERK activity in memory reconsolidation and the role of dimerization in regulating ERK subcellular signaling (Casar et al., 2008), we investigated whether ERK dimerization influences IA memory stabilization after reactivation. The small molecule DEL-22379 (DEL) has been shown to inhibit ERK dimerization without affecting its phosphorylation, and with no apparent toxicity (Herrero et al., 2015; Zaballos et al., 2022). However, ERK dimerization has not yet been characterized in the nervous system.

Consequently, our first experiment aimed to study ERK2 dimerization (dERK2) in the rodent brain (fig. 1). To induce ERK2 dimerization, two protocols were used: 1) chemical LTP (cLTP) induced in rat primary neuronal cultures using glycine stimulation (Swanger et al., 2013) and 2) intrahippocampal EGF administration, an ERK1/2 activator that also induces its dimerization (Casar et al., 2008). Next, protein-enriched samples were resolved by native PAGE. Initial validation of dERK2 assessment was performed in HEK-293T after EGF stimulation (fig. 1A) (Herrero et al., 2015). Furthermore, glycine-induced cLTP also increased dERK2, which was abolished by 30 min-incubation with DEL (fig. 1A), showing that ERK dimerization occurs in primary neuronal cultures.

**Figure 1.**
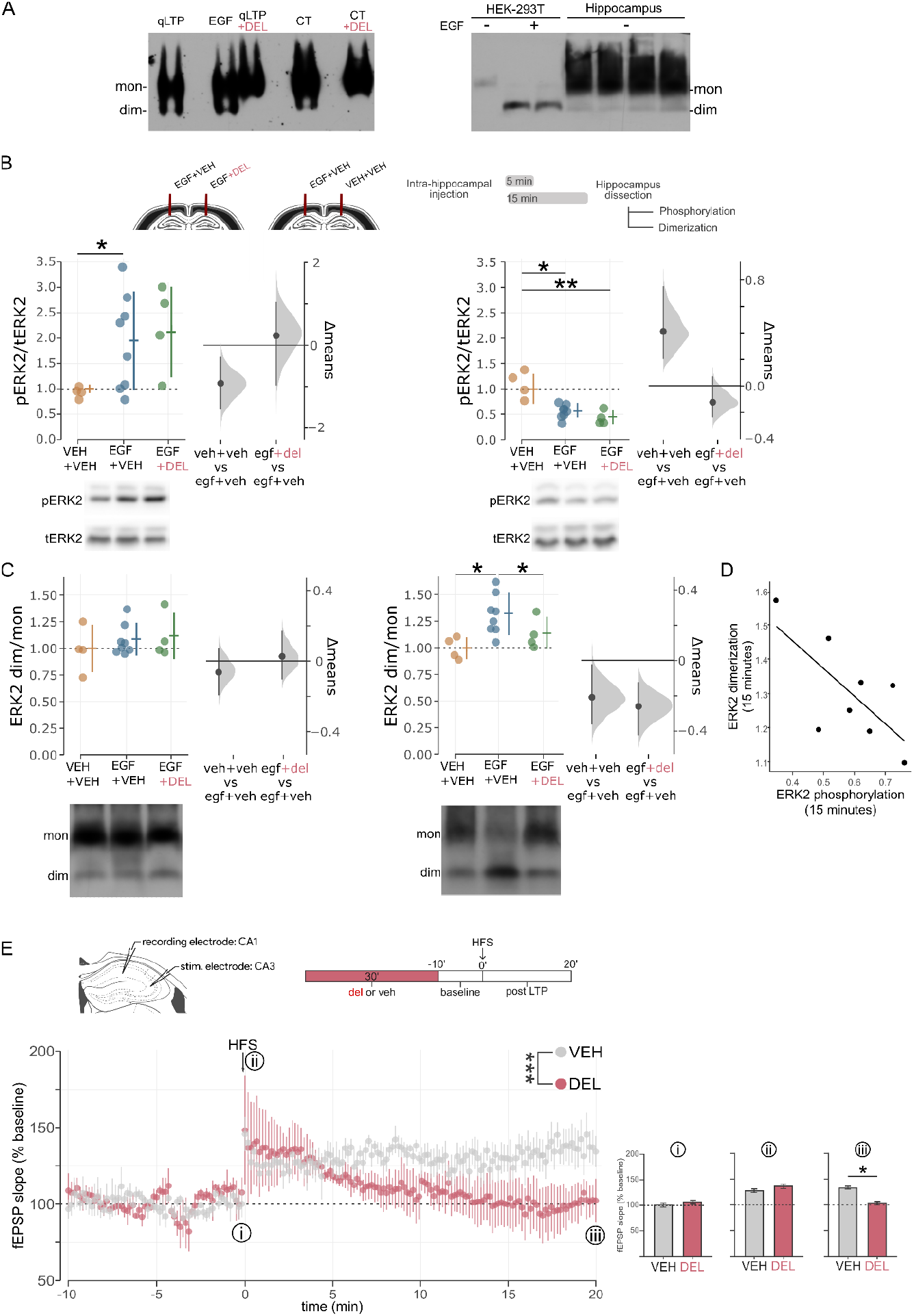
In vivo ERK2 dimerization in the rodent nervous system. **A)** Native PAGE western blot showing ERK2 dimers (dim) and monomers (mon) in samples from rat neuronal primary cultures subjected to either cLTP or EGF stimulation and-or DEL treatment (left). Control and EGF-stimulated HEK-293T culture cells and CF-1 mice hippocampal protein homogenates (right). **B-C)** ERK2 phosphorylation precedes dimerization in EGF-stimulated CF-1 mice hippocampal tissue (schematic experimental design is shown on top of fig. B; representative western blot images are shown below each graph). Hippocampal ERK2 phosphorylation **(B)** and dimerization **(C)** were assessed 5 (left) or 15 (right) min after *in vivo* local EGF plus VEH or DEL administration. Dimer / monomer (dim/mon) ratios were normalized against VEH-VEH mean value in each case. Circles represent individual mice measurements, single lines represent mean ± standard deviation (SD), and estimation plots shown on the right of each graph represent the difference between means (black dot) and the bootstrapped distribution of means (grey area) with 95% confidence intervals. **\***, p < 0.05; **\*\***, p < 0.01 in Tukey comparisons. n = 4 mice / group. **D)** Pearson correlation between ERK2 phosphorylation and dimerization assessed at 15 min in figures B and C. R^2^ = 0.71; p < 0.05. **E)** DEL pre-treatment impairs LTP in CA1 hippocampal acute slices (schematic stimulation/recording sites and experimental design is shown on top). Circles represent the mean of 4 replicates and single lines represent SD Three points of interest are indicated throughout the recording: immediately before stimulation (i), immediately after stimulation (ii), and approx. 18-20 min after stimulation (iii). Bars in i), ii) and iii) are the mean ± SD. ***, p < 0.05 (repeated measures comparison between VEH- and DEL-treated groups). n = 4 mice / group.

We next evaluated dERK2 *in vivo*. In the second experiment, we assessed the efficacy of DEL to inhibit dERK2 in CF-1 mice hippocampus. We acutely injected either PBS or EGF plus either VEH or DEL in each hemi-hippocampus of CF-1 anesthetized mice. Afterwards, mice were euthanized, and we performed SDS and native PAGE and western blot of hippocampal samples (schematic experimental design on top of fig. 1B). Consistent with previous reports (Marshall, 1995; Nguyen et al., 1993; Traverse et al., 1992; Xing and Imagawa, 1999), EGF administration increased hippocampal pERK2 levels at 5 min (fig. 1B left; F_2,13_ = 5.381, p = 0.048; VEH-VEH vs EGF-VEH: p = 0.049), followed by a significant decrease at 15 min (fig. 1B right; F_2,13_ = 8.582, p = 0.010; VEH-VEH vs EGF-VEH: p = 0.010). Treatment with DEL did not affect EGF-induced pERK2 at the time points assessed (EGF-VEH vs EGF-DEL: p = 0.989 at 5 min; p = 0.555 at 15 min), which were also statistically different from control (VEH-VEH vs DEL-EGF: p = 0.051 at 5 min; p = 0.009 at 15 min). However, dERK2 (fig. 1C) showed a distinct time course after EGF administration: it did not change at 5 min (fig. 1C left; F_2,13_ = 0.270; p = 0.769), but significantly increased at 15 min (fig. 1C right; F_2,13_ = 11.623; p = 0.004; EGF-VEH vs VEH-VEH: p = 0.014) as shown previously (Herrero et al., 2015). Administration of DEL restored dimer / monomer (dim/mon) levels to those of VEH-treated controls (EGF-VEH vs EGF-DEL: p = 0.030; VEH-VEH vs EGF-DEL: p = 0.915). Furthermore, phosphorylation and dimerization of ERK2 showed a strong negative correlation 15 min after stimulation (fig. 1D, p = 0.049). Thus, under our experimental conditions DEL prevents dERK2 induced by EGF in CF-1 mice hippocampi without affecting its phosphorylation.

To assess whether the inhibition of dERK2 has an impact on hippocampal plasticity we performed LTP experiments on hippocampal slices. Slices were incubated for 30 min with DEL or VEH and after drug washout baseline fEPSP were recorded for 10 min, followed by high frequency stimulation (HFS) on Schaffer Collaterals. While HFS was able to induce LTP in the CA1 in both DEL and VEH treated slices (immediately post-HFS: VEH = 1.45 ± 0.12, DEL = 1.47 ± 0.16, p = 0.058), fEPSPs slope in DEL-treated slices returned to baseline levels (20 min after HFS: VEH = 1.34 ± 0.21, DEL = 1.02 ± 0.20, p = 0.022), suggesting that ERK dimerization is not necessary for CA1 LTP induction, but its inhibition prevents the maintenance of HFS-induced potentiation.

These results demonstrate that DEL selectively inhibits dERK2 in the mouse hippocampus without affecting its phosphorylation and suggest that dimerization plays a critical role in hippocampal synaptic plasticity.

### Memory reactivation after Weak IA decreases hippocampal ERK dimerization

Since ERK dimerization can modulate hippocampal plasticity, and IA memory reactivation triggers cytoplasmic ERK signaling 45 min after reactivation (Krawczyk et al., 2015), we investigated whether hippocampal dERK is also regulated by memory reactivation at this time point. Animals were trained using either strong (sIA) or weak (wIA) versions of the IA task and subjected to memory reactivation (RE) 48 h later. Identically trained animals not exposed to reactivation (NR) served as control. Animals were euthanized 45 min after reactivation or at the corresponding time point for NR groups, and dERK2 was assessed in hippocampal protein samples (fig. 2).

**Figure 2.**
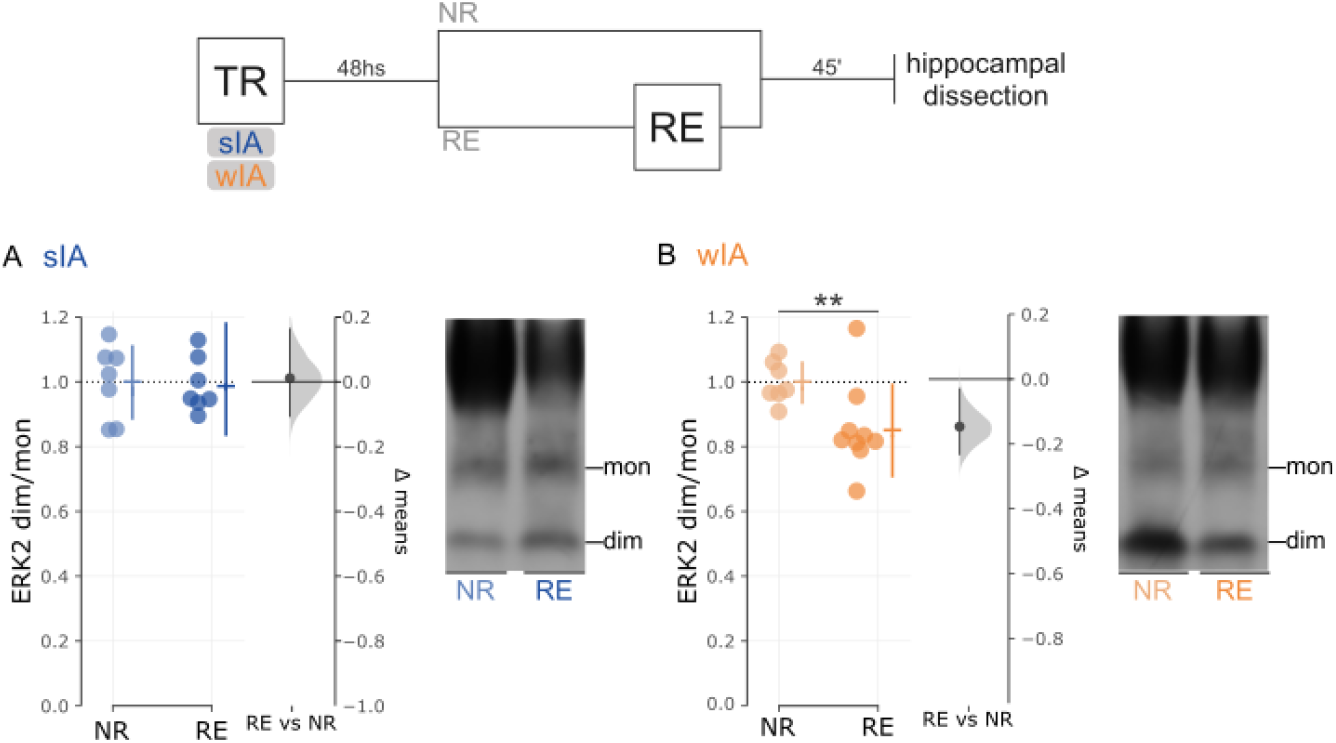
Hippocampal dERK2 after memory reactivation. **Top)** Schematic experimental design for dERK2 assessment. **Bottom)** Hippocampal dERK2 assessment by native PAGE western blots 45 after either sIA **(A)** or wIA **(B)** memory reactivation. Dim/mon ratios are normalized against NR mean value for each experiment. Color circles indicate individual values and error bars shown on the right of each set depict mean ± SD values. Estimation plots shown on the right of each graph represent the difference between means (black dot) and the bootstrapped distribution of mean difference with 95% confidence intervals. Representative western blots are also shown (right). **\***, p < 0.05; **\*\***, p < 0.01 in Tukey comparisons. n = 7-9 / group.

Reexposure to the training context 48 h after sIA did not induce significant changes in hippocampal dERK2 relative to NR controls 45 min post-reactivation (fig. 2A; *t*_13_ = 0.161; p = 0.873). In contrast, wIA memory reactivation led to a significant decrease in dERK2 compared to the NR control group (fig. 2B; *t*_14_ = 2.569; p = 0.022).

The finding that memory reactivation induced differential dERK2 depending on training strength suggests the possibility that this molecular mechanism plays a role in memory reconsolidation.

### Post-retrieval hippocampal ERK2 dimerization supports strong IA memories

Given that dERK2 is differentially regulated after memory reactivation, its inhibition might exert distinct effects on memory reconsolidation. Consequently, sIA-trained animals were bilaterally injected with either VEH or DEL in the hippocampus 35 min after memory reactivation or the corresponding time for NR groups (fig. 3A) and were euthanized 10 min later to assess hippocampal dERK2. As shown in figure 3A (lower panel), memory reactivation alone did not induce changes in dERK2 levels (F_2,21_ = 5.559; p = 0.012; NR-VEH vs RE-VEH: p = 0.336). However, hippocampal DEL injection after reactivation significantly reduced dERK2 (RE-DEL vs NR-VEH: p = 0.010), mimicking the reduction observed after wIA reactivation (fig. 2B).

**Figure 3.**
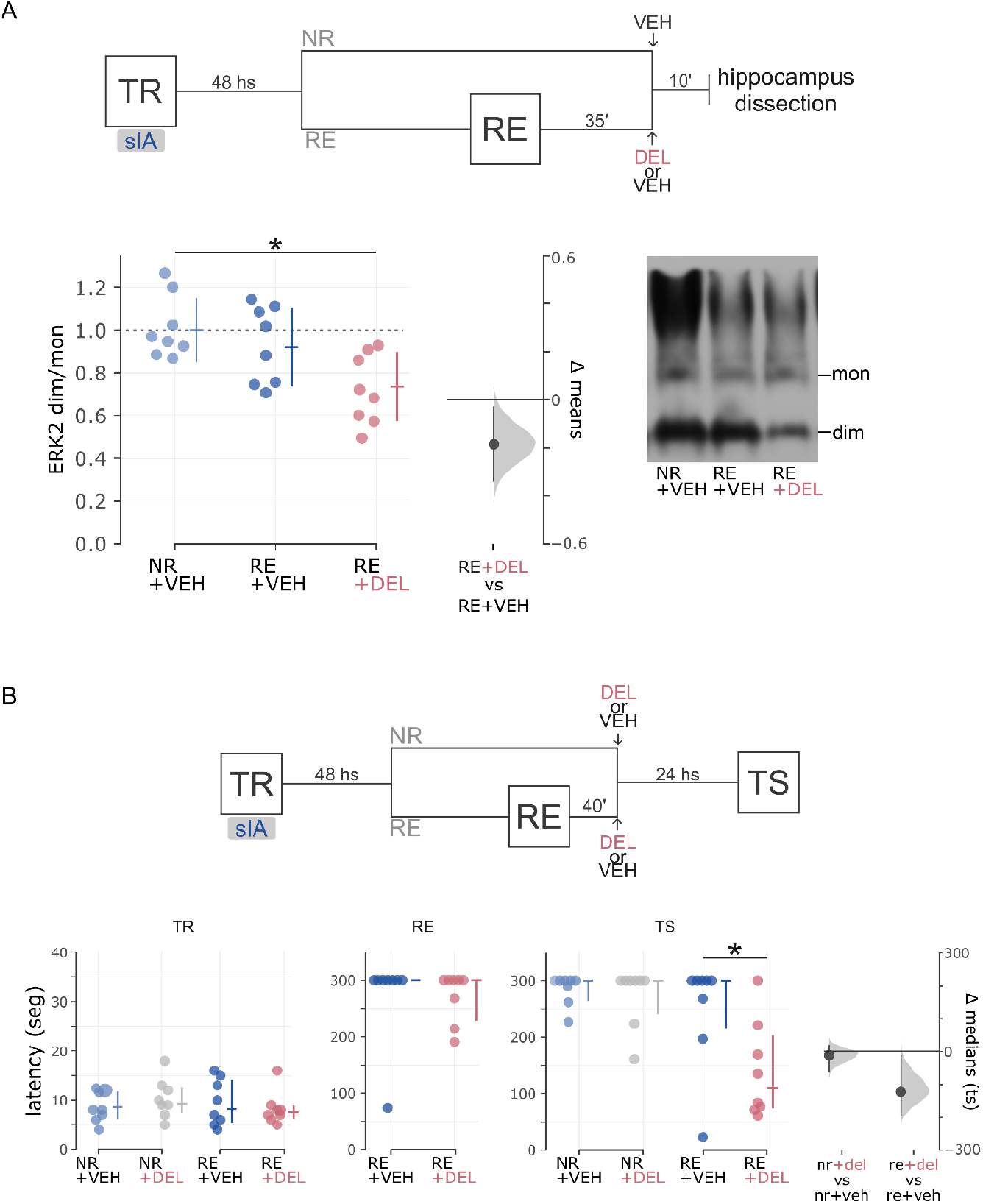
Role of hippocampal dERK2 after sIA memory reactivation. **A)** DEL decreases hippocampal dERK2 after sIA memory reactivation. Schematic experimental design for hippocampal dERK2 assessment is shown **(top)**.Hippocampal dERK2 was assessed 45 min after sIA memory reactivation and 10 min after VEH or DEL intra-hippocampal administration. Dimer to monomer ratios were normalized against NR mean value for each experiment. Color circles indicate individual values and error bars shown on the right of each dataset depict mean ± SD values. Estimation plot on the right represents the difference between means (black dot) and the bootstrapped distribution of mean difference with 95% confidence intervals. Representative western blots are also shown (right). **\***, p < 0.05; **\*\***, p < 0.01 in Tukey adjusted *post hoc* comparisons. n = 8 / group. **B)** Inhibition of hippocampal dERK2 after strong IA memory reactivation impairs LTM. Schematic experimental design is shown **(top)**. Color circles indicate individual latencies to step through values and error bars shown on the right of each set depict median ± inter quartile range (IQR). Estimation plot on the right of TS graph represents the difference between means (black dot) and the bootstrapped distribution of means (grey area) with 95% confidence intervals. **\***, p < 0.05; **\*\***, p < 0.01 in Tukey adjusted *post hoc* comparisons. n = 8 / group.

To determine whether basal dERK2 levels are necessary for strong IA memory reconsolidation, we administered either VEH or DEL 40 min after strong IA memory reactivation and tested retention 24 h later (fig. 3B). A significant interaction between reactivation and drug treatment was observed on latencies during TS (F_2,28_= 5.291; p = 0.029). DEL-treated RE animals exhibited significantly reduced latencies compared to VEH-treated ones (RE+VEH vs RE+DEL: p = 0.029), whereas latencies in VEH- and DEL-injected NR groups were not statistically different (NR+VEH vs NR+DEL: p = 0.983). Furthermore, latencies in VEH-injected groups were consistent across conditions (RE-VEH vs NR-VEH: p = 0.685). This impairment was sustained for at least a week (suppl. fig. 2), suggesting a specific effect on the reconsolidation process (Quirk, 2002; Rescorla, 2004).

Together, these findings indicate that basal dERK2 is critical for the reconsolidation of sIA memory.

### Inhibiting hippocampal ERK dimerization after reactivation strengthens weak IA memories

We next investigated whether further inhibiting dERK2 after wIA reactivation would affect memory reconsolidation by training four groups of animals in a wIA task, two of which were submitted to memory reactivation (RE) 48 h later. One NR and one RE group of animals were administered VEH, while the remaining groups were injected DEL in the dorsal hippocampus 40 min after RE (fig. 4A). Testing 24 h later revealed a significant interaction between reactivation and drug treatment (F_2,30_ = 13.254; p = 0.001). Remarkably, DEL-treated RE animals showed significantly higher latencies than the VEH-treated ones (p = 0.005). This enhancement was specific to memory reactivation, as animals from NR-VEH and NR-DEL groups did not perform significantly different during the test session (p = 0.379). Furthermore, VEH-treated groups did not differ statistically during the test session (p = 0.273).

**Figure 4.**
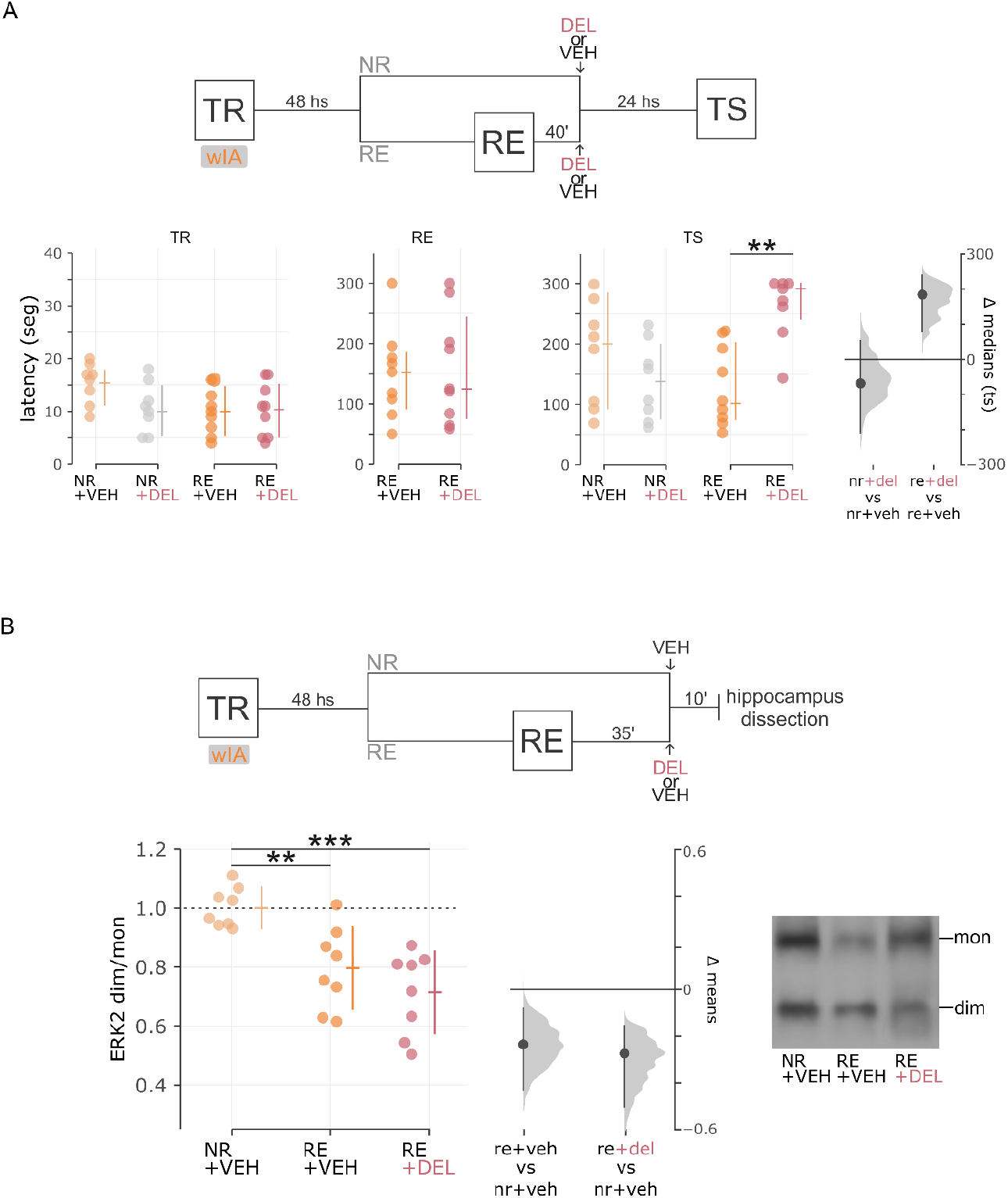
Role of hippocampal dERK2 after wIA memory reactivation. **A)** Inhibition of hippocampal dERK2 after strong IA memory reactivation improves LTM. Schematic experimental design is shown on top. Color circles indicate individual latencies to step through values during training (TR, left), reactivation (RE, center) and testing (TS, right). Error bars shown on the right of each dataset depict median ± IQR. Estimation plot on the right of TS graph represents the difference between means (black dot) and the bootstrapped distribution of mean difference with 95% confidence intervals. **\***, p < 0.05; **\*\***, p < 0.01 in Tukey adjusted *post hoc* comparisons. n = 8 - 9 / group. **B)** Hippocampal dERK2 decreases after wIA memory reactivation. Schematic experimental design is shown on top. Color circles indicate individual values and error bars shown on the right of each dataset depict mean ± SD values. Estimation plots on the right of the graph represent the differences between means (black dot) and the bootstrapped distribution of means (grey area) with 95% confidence intervals. Representative western blots are also shown (right). **\***, p < 0.05; **\*\***, p < 0.01 in Tukey adjusted *post hoc* comparisons. n = 8 / group.

Not only did DEL administration improve retention in RE groups, but it also increased the proportion of animals with higher latencies during TS compared to RE (suppl. fig. 3). A significant positive correlation was observed between RE and TS response levels in DEL-RE animals (R^2^ = 0.73; p = 0.025), contrasting with the VEH-RE group (R^2^ = 0.42; p = 0.270).

Finally, dERK2 levels were significantly modulated by reactivation in VEH- and DEL-injected animals (fig. 4b, F_2,21_ = 13.254; p = 0.0002). In contrast to what we observed after sIA training, wIA memory reactivation led to a significant decrease in dERK2 (see fig. 2B) which was also observed after VEH-administration in mice hippocampi (fig. 4B; NR+VEH vs RE+VEH: p = 0.006). However, DEL administration did not significantly affect dERK2 levels following memory reactivation (RE+VEH vs RE+DEL: p = 0.375), though these levels were significantly different from those in the control group NR-VEH (NR+VEH vs RE+DEL: p = 0.0003).

These results demonstrate that inhibiting dERK2 after reactivation enhances retention of wIA memories, suggesting that the decrease in dERK2 associated with wIA reactivation may represent a boundary condition for memory reconsolidation that can be shifted by pharmacological intervention.

## DISCUSSION

In this work, we show for the first time that ERK dimerization constitutes a key regulatory step in neuronal plasticity and memory processes. Several lines of evidence indicate that subcellular localization of a particular intracellular signaling pathway activity is crucial for specific biological outcomes in different systems (Afinanisa et al., 2021; Casar and Crespo, 2016; Formoso et al., 2020), including the central nervous system (Salles et al., 2015). Over the past decade, oncological research has advanced the development of drugs targeting key pathways to reverse pathological conditions. Interestingly, many of these pathways have also been implicated in learning and memory processes. However, their regulatory mechanisms within the nervous system often remain poorly understood. ERK signaling serves as a notable example. ERK phosphorylation is well known to be essential for neural plasticity and cognitive processes, and its deregulation often leads to pathological conditions (reviewed in Ojea Ramos et al., 2022). ERK dimerization has been proposed to regulate the subcellular location of its activity, thereby influencing the phosphorylation of specific targets and ultimately the cellular fate (Casar and Crespo, 2016; Herrero et al., 2015). However, despite extensive studies on neural ERK activation, the functional role of ERK dimerization in the nervous system remains unexplored.

We first established that ERK dimerization can be stimulated in neural tissue, both *in vitro* using primary neuronal cultures and *in vivo* within the hippocampus of CF-1 mice, consistent with previous studies in EGF-stimulated cell lines (Herrero et al., 2015). To validate our findings, we replicated Herrero and colleagues’ experiments as positive control for dERK2 assessment in rodent neural tissue (fig. 1A). Like many other tissues, the brain expresses EGF receptors, which regulate neurite outgrowth and neuronal homeostasis (Romano and Bucci, 2020). However, distinct ligands can induce different ERK activation kinetics and cellular outcomes (Miningou and Blackwell, 2020). Recent research on breast tumors has demonstrated that ligand-dependent ERK dimerization is crucial for cellular motility as its disruption impairs actin cytoskeleton rearrangements (de la Fuente-Vivas et al., 2024). In physiological contexts, EGF concentration partially determines cellular responses, influencing the balance between migratory and proliferative states (e.g. during early wound closure, low EGF concentrations promote pro-migratory signaling at the plasma membrane). How ERK dimerization is modulated by ligand specificity and concentration, as well as its functional consequences in neurons remain open questions for future investigation.

### ERK dimerization in plasticity

Hippocampal plasticity is linked to memory formation, with ERK signaling playing a crucial role in LTP. Early studies showed that HFS induces ERK2 phosphorylation in the CA1 of rat hippocampus, and blocking ERK phosphorylation before LTP induction prevents long-term maintenance (English and Sweatt, 1997). Later, multiple studies have provided evidence supporting its role in these processes (Atkins et al., 1998; Impey et al., 1998; Kelleher et al., 2004; Selcher et al., 2003). Building on this, we hypothesize that ERK dimerization regulates synaptic plasticity and memory. We found that DEL pre-treatment impaired long-term LTP stabilization without affecting its induction. These results suggest that both ERK phosphorylation and dimerization are key for LTP maintenance. Furthermore, ERK signaling is compartmentalized within subcellular regions (Casar and Crespo, 2016), indicating that dimerization may regulate distinct ERK pools contributing to plasticity and memory. Since ERK dimerization and phosphorylation target different compartments, multiple pathways are likely triggered during ERK signaling. Memory processes activate various receptor systems that influence ERK signaling kinetics. Thus, ERK dimerization likely plays a role in LTP induction, though the precise dynamics remain unclear.

### ERK dimerization in inhibitory avoidance memory reconsolidation

To this day, ERK signaling has been considered a key pathway for LTM consolidation (Atkins et al., 1998; Blum et al., 1999; Feld et al., 2005; Schafe et al., 2000; Selcher et al., 2003; Walz et al., 1999), labilization and reconsolidation (Krawczyk et al., 2016, 2015; Nagai et al., 2007; Raut et al., 2024); and has been shown to become activated in the same set of neurons engaged during conditioning upon memory reactivation (Zamorano et al., 2018). It is worth noting that ERK’s role in plasticity and memory processes has been attributed almost exclusively to phosphorylation, leaving the potential involvement of other post-translational modifications in these processes largely unexplored. In this work, our focus was on reconsolidation mechanisms assessed 48 h post-training. However, we acknowledge that earlier time points (e.g., 30–45 min after training) should be examined to determine whether ERK2 dimerization also contributes to the initial consolidation. This remains an open question for future studies.

Our results reveal a dual effect of DEL administration on IA memory reconsolidation, depending on the strength of the stimuli used during TR. After sIA TR, intra-hippocampal DEL administration impaired memory reconsolidation, while after wIA TR it enhanced memory retention. These findings suggest that the molecular mechanisms engaged upon memory reactivation are dependent on TR conditions, with hippocampal ERK dimerization playing opposite roles in the behavioral outcome of sIA and wIA reactivation. Notably, it has been reported that pharmacological inhibition of ERK dimerization increases its nuclear signaling (Casar et al., 2008). In this line, sIA and wIA may engage different spatial ERK pools that could explain the bidirectional effect observed, probably involving different downstream cytoplasmic targets such as MNK or RSK, depending on the training intensity. Although we did not directly assess ERK2-protein interactions, future studies could elucidate whether different reactivation conditions alter ERK2 binding partners. Moreover, distinct pool-specific components (e.g. upstream ligands and/or downstream targets) of the ERK pathway may guide different cellular fates (Pirbhoy et al., 2017). Furthermore, different Ras isoforms have recently been shown to induce opposing synaptic plasticity processes (López-Merino et al., 2025). Considering the finding that BRAF- and RAS-mutant thyroid cells respond differentially to DEL (Zaballos et al., 2022), the possibility of different mechanisms of action of the inhibitor on memory processes cannot be excluded. On the other hand, sIA and wIA memory reactivation might recruit distinct neural circuits within the hippocampus as it was observed that protein synthesis inhibition after reactivation exerted differential effects on memory restabilization of an IA task in mice according to the area infused (Fukushima et al., 2014). Further experiments will be needed to elucidate how ERK dimerization exerts a bidirectional effect on memory reconsolidation, depending on the strength of the training stimuli.

Our experiments suggest that the effect on hippocampal ERK dimerization is specific to the reconsolidation process, as non-reactivated (NR) animals did not show changes in step through latency following DEL treatment (figures 3B and 4A). Furthermore, sIA reconsolidation memory impairment following DEL administration persisted for at least one week (suppl. fig. 2), suggesting that there was no spontaneous recovery of the original memory. However, the possibility of the enhancement of an extinction process cannot be completely ruled out and future work using reinstatement or renewal paradigms would help to distinguish between these mechanisms. These findings may have therapeutic relevance, particularly for conditions such as post-traumatic stress disorders or phobias, in which reconsolidation-based interventions are being actively explored. In contrast to conventional ERK inhibitors, targeting ERK dimerization could allow for more specific modulation of memory processes without broadly suppressing plasticity mechanisms. Based on previous work (Herrero et al., 2015), we employed a single DEL dose and specifically assessed whether it impaired ERK dimerization in neural tissue (fig. 1); further pharmacological profiling, including dose-effect studies, would help clarify the compound’s potency and specificity. It is important to note that only male mice were used in this study. Given known sex differences in ERK signaling and memory processes, future work should examine whether these findings generalize to female subjects. Also, although DEL reduced ERK2 dimerization in vitro and in vivo, its behavioral effect in the wIA group—where dimerization is already inhibited—suggests possible transient or subcompartmental actions not captured by our biochemical assays.

In the present manuscript, we provide the first evidence of ERK2 dimerization playing a critical role in memory and plasticity processes, particularly in the context of hippocampal plasticity and memory reconsolidation, thus introducing a novel regulatory mechanism in the brain. Future studies are needed to elucidate the other molecular players involved in this mechanism.

## Supporting information

Supplemental figures 1 - 3 with legends

## Abbreviations

ERK1/2: extracellular signal-regulated kinase 1/2
IA: Inhibitory avoidance
LTM: long-term memory
LTP: long-term potentiation
wIA: weak Inhibitory Avoidance
sIA: strong Inhibitory Avoidance
PFA: paraformaldehyde
RE: memory reactivated animals
NR: non-reactivated animals
EGF: epidermal growth factor
DEL: DEL-22379
Veh: Vehicle
DMSO: dimethyl sulfoxide
IQR: Interquartile range
TR: training
RE: memory reactivated animals
TS: testing

## Author Contributions

S.O.R. performed experiments, analyzed data, and contributed to writing parts of the manuscript. C.M. performed behavioral experiments, analyzed data, and reviewed the manuscript. M.C.K., J.M. and M.F.A. performed experiments and reviewed the manuscript. M.V.B. performed experiments and contributed to writing parts of the manuscript. M.M.B contributed to designing the experiments, performed behavioral experiments and interpreted data. A.G.R contributed to data analysis and reviewed the manuscript. F.J.U. & S.O.R. designed and performed electrophysiology experiments in slices, interpreted data. M.F. proposed the original research project, designed experiments and performed biochemical assays, interpreted data, drafted the manuscript and is the corresponding author.

## Funding

This work was supported by the following grants: Agencia Nacional de Promoción de la Investigación, el Desarrollo Tecnológico y la Innovación (ANPCYT PICT 2016 0295, ANPCYT PICT 2020 01534 and PICT 2015 1199), Consejo Nacional de Investigaciones Científicas y Técnicas (CONICET PIP 2014-2016 No. 11220130100519CO) and Universidad de Buenos Aires (UBACYT 2018-2021 - 20020170100390BA and 2014-2017 - 20020130200283BA), Argentina.

## Competing Interests

The authors have no competing interest to disclose.

## Declaration of generative AI and AI-assisted technologies in the writing process

During the preparation of this work the author(s) used ChatGPT in order to improve language and readability of the manuscript. After using this tool/service, the first and corresponding authors reviewed and edited the content as needed, and take full responsibility for the content of the publication.

## Notes

### Competing Interest Statement

The authors have declared no competing interest.

### Summary of Updates

Some aspects of the discussion were revised in order to clarify.

