## Supplemental figures 1 - 3 with legends for "Hippocampal ERK2 dimerization is critical for memory reconsolidation and synaptic plasticity"

### Supplementary figures and legends

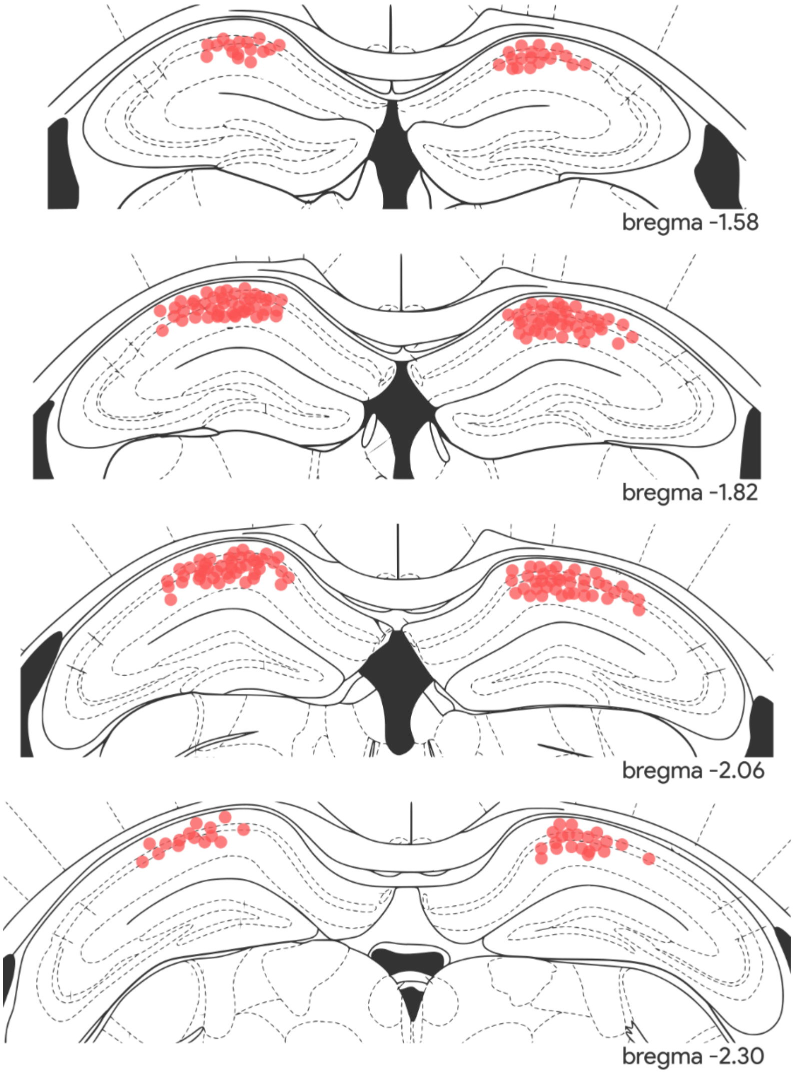


**Supplementary figure 1:** Injection site locations for all the animals used in the experiments presented in this study.

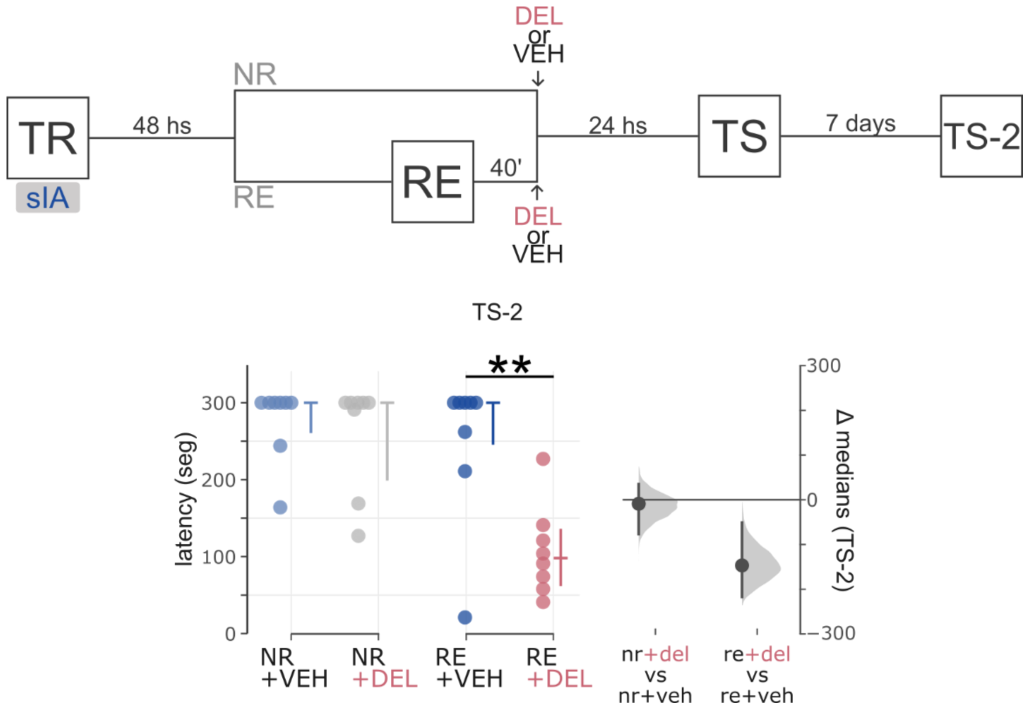


**Supplementary figure 2:** Inhibition of hippocampal dERK2 after sIA memory reactivation impairs LTM persistently (TS-2 performed 7 days after TS in fig. 3B). Schematic experimental design is shown in top panel. Color circles indicate individual latencies to step through values and error bars shown on the right of each dataset depict median ± IQ intervals. Estimation plots shown on the right represent the difference between medians (black dot) and the bootstrapped distribution of medians (grey area) with 95% confidence intervals. ******, p<0.01 in Tukey adjusted *post hoc* comparisons.

Statistical analysis of suppl fig. 1:
There was a significant interaction between reactivation and drug treatment on latencies during the TS-2 session ($F_{2,28}$=6.342; p=0.012), with significant main effects of reactivation protocol ($F_{1,28}$=9.723; p=0.004) and drug treatment ($F_{1,28}$=12.811; p=0.001). Post-hoc analysis revealed significant differences between VEH- and DEL-injected RE groups (p=0.001), but no changes either between NR groups (p=0.966) or between NR and RE VEH-injected animals (p=0.842).

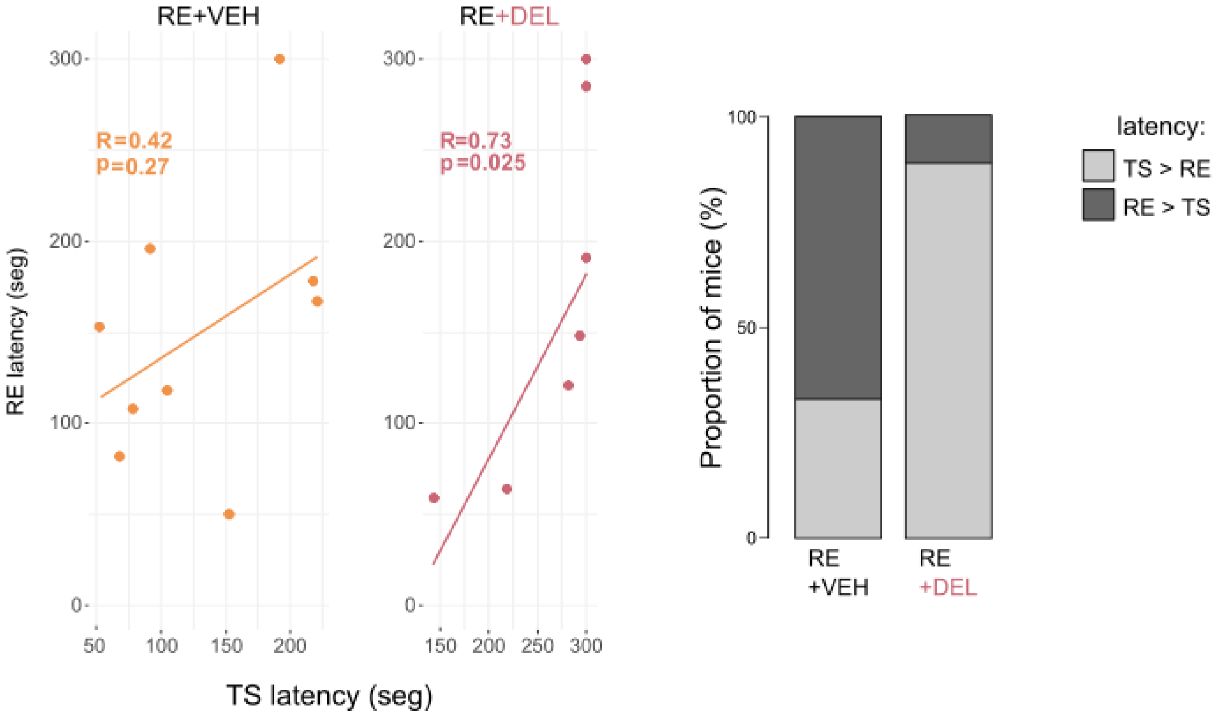


**Supplementary figure 3:** Inhibition of dERK following wIA memory reactivation improves LTM. **Left)** Pearson correlation between RE and TS latencies from VEH-RE and DEL-RE groups depicted in figure 4A. **Right)** The proportion of animals that expressed higher step-through latencies during RE vs TS for VEH- and DEL-injected RE-groups is also represented.
